# Bacteriophage host prediction using a genome language model

**DOI:** 10.64898/2026.03.19.712863

**Authors:** Zhijie Wang, Javier Arsuaga

## Abstract

Computational bacteriophage host prediction from genomic sequences remains challenging because host range depends on diverse, rapidly evolving genomic determinants—from receptor-binding proteins to anti-defense systems and downstream infection compatibility—and because the signals available to predictors, including sequence homology, CRISPR spacer matches, nucleotide composition, and mobile genetic elements, are sparse, unevenly distributed across taxa, and constrained by incomplete host annotations. Here, we frame host prediction as an unsupervised retrieval problem. We asked whether embeddings from the pretrained genome language model Evo2 captured a reliable host-range signal without training on phage–host labels. We generated whole-genome embeddings for phages and candidate bacterial hosts with the Evo2-7B model, applied normalization, and ranked hosts by cosine similarity. Using the Virus-Host Database, we selected embedding and fusion choices on a Gram-positive validation cohort and then evaluated the approach on a held-out Gram-negative test cohort to minimize data leakage. We found that Evo2 was strongest at retrieving multiple plausible hosts, with the recorded host in the top 10 for 55.4% of phages. However, it did not maximize species-level top-1 accuracy (19.4% vs. 23.2% for the best baseline). At higher taxonomic ranks, Evo2 captured a coarser host-range signal: top-1 accuracy reached 43.4% at the genus level and 51.6% at the family level. Reciprocal rank fusion of Evo2 with BLASTN, VirHostMatcher, and PHIST improved all retrieval metrics. Top-10 retrieval rose to 58.5% and top-1 accuracy to 26.9%. Stratified analyses by phage genome length, host clade, and host mobile genetic element coverage revealed scenario-dependent performance. Evo2 embeddings excelled for intermediate-length phages and when host mobile element content was low, whereas alignment and *k*-mer methods dominated when local homology was abundant. These results suggest that pretrained genome embeddings complement established alignment- and k-mer/composition-based methods and that context-aware hybrid pipelines may help improve phage host prediction.

**Author summary:** Bacteriophages are viruses that prey on bacteria and play central roles in microbial ecosystems, nutrient cycling, and the spread of antibiotic resistance genes. Knowing which bacterium a phage can infect is important for applications such as phage therapy, where viruses are used to treat bacterial infections, but making this prediction from DNA sequence data alone remains difficult. Existing computational tools each exploit different types of genomic evidence, and none works reliably across all settings. We asked whether an artificial intelligence model trained to read raw DNA—without ever being shown which phages infect which hosts—could contribute a new, complementary signal. We found that this approach was particularly effective at narrowing the field to a short list of candidate hosts and at capturing broad evolutionary relationships between phages and bacteria. When we combined it with established sequence-comparison tools, overall prediction improved beyond what any single method achieved alone. By examining when each method succeeded or failed, we identified biological factors that govern prediction difficulty, offering practical guidance for building more robust prediction systems.

## Introduction

Bacteriophages (phages) are an essential component of microbiomes, including those found in soils, oceans and the human body [1–6]. They shape community structure and influence microbial ecology and evolution by propagating in different bacterial communities through lysogenic and lytic pathways or as independent plasmids [7, 8].

Identification of phage hosts remains a challenging problem. Wet-lab assays remain the gold standard for validating phage–host interactions. Nonetheless, these assays can be slow, costly, and biased toward culturable systems [7, 9, 10]. Some of these challenges have been overcome by metagenomic studies; these, however still lack the necessary information to unequivocally associate a phage and its host [7, 9, 10]. These challenges make sequence-based phage host prediction a central computational problem.

Existing computational approaches exploit different genomic signals. Some methods use direct evidence, such as exact DNA matches between bacterial and viral genomes. Examples include CRISPR spacer hits [11], alignment-based methods [12] and *k*-mer frequency profiles [13–17]. These methods, however, are not fully reliable because broad genomic similarity does not necessarily translate into shared host range since infection specificity can be modular and concentrated in a small number of loci (e.g., tail fibers and receptor-binding proteins) that diversify quickly [18–20]. For instance, alignment approaches can work well when local homology exists, but this signal may be absent or altered in novel lytic phages [9]. Composition-based methods, on the other hand, leverage host-like oligonucleotide usage, but this signal is indirect, and can be confounded by GC content, shared ancestry, cohabitation in similar environments, and mosaic genome architecture [9, 21, 22]. Overall performance can also degrade for short genomes or fragmented assemblies [22, 23].

More recently supervised or unsupervised machine learning models have been proposed [23–31]. Genome-level approaches learn whole-sequence representations from nucleotide sequences [26, 27, 29, 30]; protein-level work often embeds receptor-binding proteins with pretrained protein language models [19, 20]; hybrid approaches also combine DNA and protein sequence features [31]. These approaches can outperform single-signal methods when host information is retained in genome-wide or receptor-binding-protein features and can be integrated by the model [20, 22, 23, 27]. However, most of these approaches require known interactions or host labels for training, which can limit their ability to generalize to under-sampled host taxa or novel phage lineages [22, 23]. Protein-centric methods rely on accurate identification of receptor-binding proteins or other functional annotations, which are often incomplete in fragmented or poorly characterized genomes [18, 20]. Taken together, each class of method — alignment, composition, and supervised learning — captures a different facet of host signal, yet none is reliable across all settings.

Recent genome foundation models, including `Evo2`, learn general-purpose representations directly from DNA sequences [32, 33]. Without task-specific fine-tuning, Evo2 representations capture biologically meaningful features. For instance, sparse-autoencoder analyses have been shown to recover prophage-associated features and transcription-factor binding motifs, while supervised models trained on frozen embeddings achieve strong performance on exon classification and BRCA1 variant classification [32]. These findings suggest that `Evo2` embeddings encode rich functional information across diverse genomic contexts.

A natural question for host prediction is therefore whether pretrained genome embeddings encode host-range signals without training on phage–host labels. In this work, we framed phage–host prediction as an unsupervised retrieval problem. For each query phage, we produced a ranked list of putative hosts by cosine similarity in `Evo2` embedding space. We also generated host rankings using four established unsupervised baselines (`BLASTN`, `PHIST`, `VirHostMatcher`, and `WIsH`) [12–15].

Because no single signal (or method) dominates across scenarios, and a single aggregate benchmark score can hide systematic blind spots, rank fusion approaches can integrate complementary evidence without additional training [7, 9, 10, 22, 23]. We propose a reciprocal rank fusion (RRF) to produce a single ranked host list per phage that combines the ranked lists generated by these different methods.

We evaluated our approach on two cohorts from the Virus-Host Database (Virus-Host DB) [34]. These cohorts comprise phages infecting Gram-negative bacteria and phages infecting Gram-positive bacteria. Because cross-Gram infection appears rare in reported host ranges, we treated Gram-positive and Gram-negative host sets as largely disjoint [35, 36]. We used the Gram-positive cohort as a validation set to choose `Evo2` embedding extraction details and the fusion protocol. We then fixed all choices and evaluated the Gram-negative cohort as a held-out test set.

Since the Virus-Host DB has a long-tailed host distribution, we evaluated host-balanced MRR and Hit@k. These metrics prevent common hosts from dominating method performance [22]. Species-level evaluation may underestimate prediction accuracy because phage–host databases typically record only a single host association per phage, so a prediction of a closely related but unlisted species is scored as incorrect even when it may be biologically plausible. We therefore also reported Hit@1 at the genus and family levels.

On the held-out Gram-negative test cohort, `Evo2` alone was strongest at retrieving multiple plausible hosts but did not maximize species-level top-1 accuracy. At genus and family levels, `Evo2` captured host-range signal more effectively. Reciprocal rank fusion with `BLASTN`, `VirHostMatcher`, and `PHIST` improved all ranking metrics beyond any single method. We went beyond overall benchmark averages by analyzing scenario-dependent performance, asking under which conditions each signal is reliable and where it breaks down. More specifically, we stratified performance by phage genome length, host clade, and host mobile genetic element burden. This stratified analysis helps explain tool disagreement and further motivates hybrid pipelines that adapt to the biological context of each prediction.

In summary, in this work we show that frozen `Evo2` genome embeddings, combined with reference-set normalization, provide an effective signal for phage–host prediction without training on phage–host labels. We further show that label-free reciprocal rank fusion with standard alignment and *k*-mer methods improves ranking on a held-out host space. Finally, we provide scenario diagnostics—stratified by phage length, host clade, and host mobile element burden—that help explain the dominance of each genomic signal.

## Results

### Pipeline overview

Fig 1 outlines the proposed workflow. We treated host prediction as a retrieval problem in which each query phage was ranked against candidate host species. We constructed two non-overlapping cohorts from Virus-Host DB [34]: a Gram-positive cohort (3,514 phages, 209 host species) used for validation and a Gram-negative cohort (4,490 phages, 308 host species) reserved for a held-out test set to reduce data leakage during model selection (Fig 1A).

**Fig 1.**
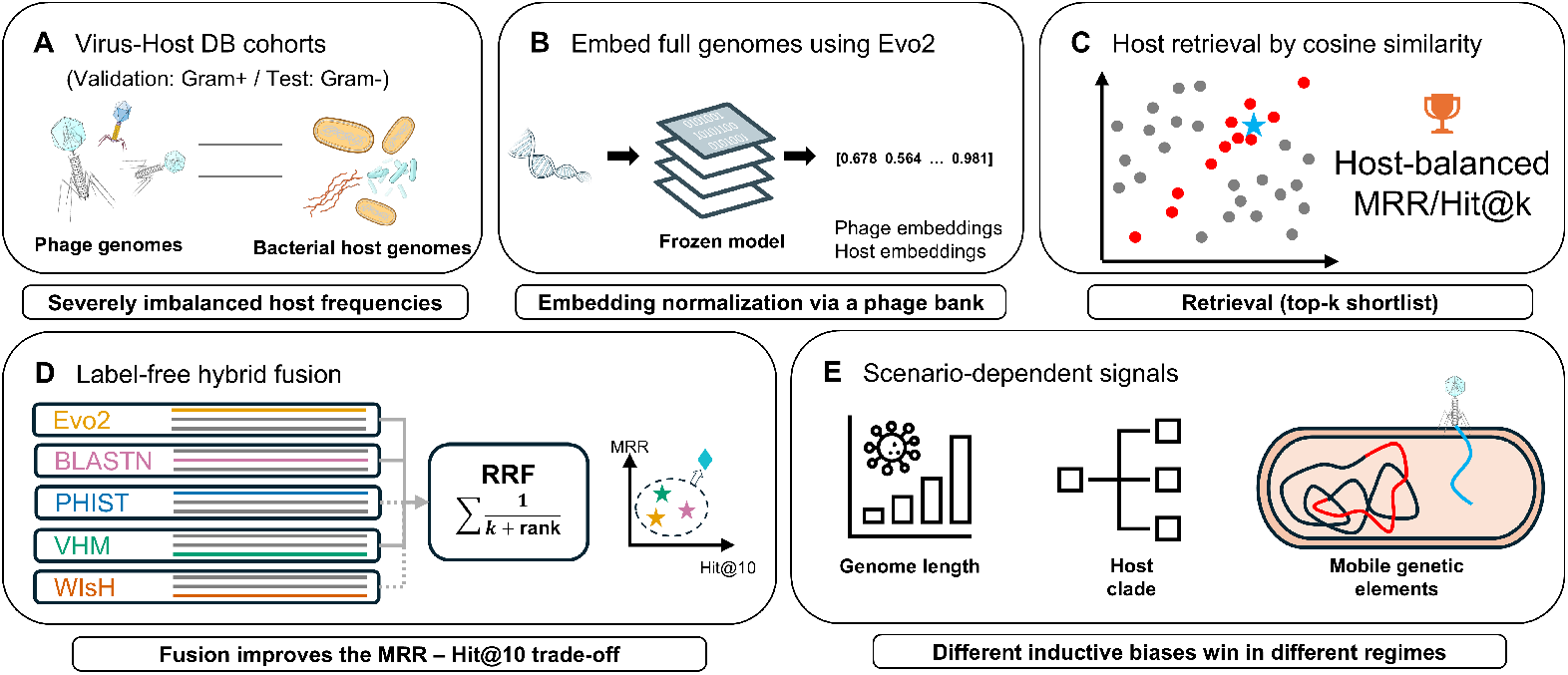
Pipeline overview. We frame phage–host prediction as an unsupervised retrieval problem over a cohort-specific host database. (A) Gram-positive (validation) and Gram-negative (held-out testing) cohorts from Virus-Host DB. (B) Whole genome embedding with frozen `Evo2-7B`. (C) Hosts ranking by cosine similarity, evaluated with host-balanced metrics. (D) Reciprocal rank fusion of `Evo2` with unsupervised baselines. (E) Stratified performance analyses.

For each phage and host genome, we extracted whole-genome embeddings from the frozen `Evo2-7B` model by mean-pooling hidden states from an intermediate block and applied normalization (Fig 1B). Candidate hosts were ranked by cosine similarity and evaluated with host-balanced retrieval metrics (Fig 1C). In parallel, we generated host rankings with four baselines that capture complementary genomic signals: `BLASTN` [12] (local sequence alignment), `VirHostMatcher` [13] (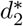 oligonucleotide composition), `PHIST` [15] (exact *k*-mer matches), and `WIsH` [14] (Markov-chain likelihood). Each baseline produced its own host ranking, and we used reciprocal rank fusion (RRF) to merge complementary rankings into a single consensus list per query phage (Fig 1D).

Finally, because no single method dominated across all scenarios, we stratified performance by phage genome length, host clade, and host mobile genetic element burden to diagnose when each genomic signal is most informative (Fig 1E). The following subsections detail each component of this pipeline.

### Virus-Host Database exhibits host-frequency imbalance

In the Virus-Host DB, a small number of bacterial host species accounts for hundreds to thousands of recorded phage–host records. For example, *Escherichia coli* is associated with 1,199 phages in the Gram-negative cohort, and *Gordonia terrae* with 534 interactions in the Gram-positive cohort. Fig 2 illustrates this imbalanced distribution and most likely reflects uneven sampling across host taxa. In order to avoid disproportionately rewarding methods that retrieved high-frequency hosts, we report host-balanced retrieval metrics, MRR and Hit@k, that down-weight queries from common hosts by inverse host frequency (see Methods).

**Fig 2.**
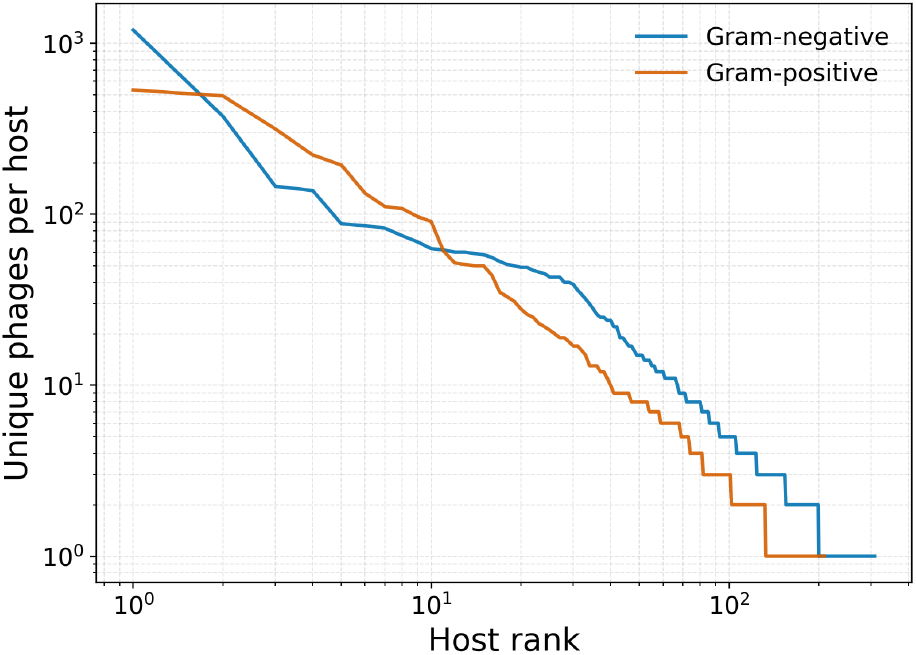
Long-tailed host frequency. Hosts are ranked by the number of associated phages, and the number of unique phages per host is plotted on log–log axes for the Gram-positive (red) and Gram-negative (blue) cohorts. A small number of hosts dominates the recorded interactions, motivating host-balanced evaluation.

### Embedding choice and normalization for optimal Evo2 host retrieval

To optimize phage–host prediction with `Evo2`-generated embeddings, we needed to select the most informative representation produced by the `Evo2` blocks and an appropriate normalization procedure for that representation. We made both choices on the Gram-positive cohort and fixed them before evaluating on the held-out Gram-negative test cohort.

`Evo2` uses a StripedHyena 2 architecture comprising 32 blocks that interleave Hyena convolutional operators (SE, MR, LI) with Transformer blocks [32]. Because each block type captures different aspects of the input sequence, the most informative layer for a given downstream task need not be the final one. In `Evo2-7B`, the final block is block 31, but Brixi et al. selected block 26, a Hyena-MR block, for sparse-autoencoder interpretability and exon–intron classification, and block 27 for the embedding-based DART-Eval variant-effect task. For supervised *BRCA1* variant classification with `Evo2-40B`, they selected block 20 [32]. We therefore evaluated sequence representations from different `Evo2` blocks to select the best block for genome representation.

We subdivided each genome into overlapping fragments of 8,192 bp and extracted representations from each fragment using a frozen `Evo2-7B` model. The `Evo2` representation for each fragment was trimmed to avoid leakage, and the retained token embeddings were then mean-pooled (Fig S1). Holding this extraction procedure fixed, we swept blocks 20–31 and observed peak retrieval performances in intermediate layers (blocks 24–27; Fig 3). Block 24 achieved the highest host-balanced MRR and was used for all downstream embeddings and results.

**Fig 3.**
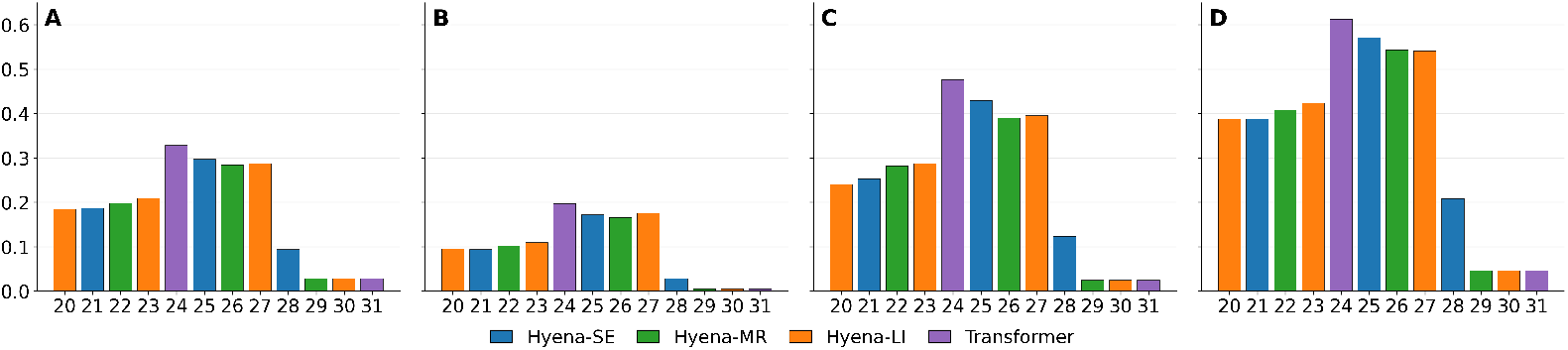
Evo2 block sweep for host retrieval. Panels show host-balanced (A) mean reciprocal rank (MRR), (B) Hit@1, (C) Hit@5, (D) Hit@10 on the Gram-positive validation cohort as a function of the `Evo2` block used to construct pooled whole-genome embeddings (blocks 20–31). Candidate hosts are ranked by cosine similarity in embedding space. Intermediate blocks (24–27) perform best, motivating the use of block 24 in downstream analyses.

We compared retrieval performance between raw embeddings and reference-set normalized embeddings. Normalization was implemented by first computing a *z*-score transform using statistics estimated from a separate reference set, and then applying *L*_2_ normalization (Fig S2; see Methods). We tested three reference-set normalization strategies on the Gram-positive cohort and found that using an external phage bank yielded the best retrieval performance (Table S1). This strategy was fixed for all subsequent analyses.

### Evo2 improves top-k host retrieval and genus/family prediction on the Gram-negative test cohort

We benchmarked host prediction methods on Gram-negative pairs from Virus-Host DB using five unsupervised methods (`BLASTN`, `PHIST`, `VirHostMatcher`, `WIsH`, and `Evo2` embeddings), none of which were trained on phage–host labels. Table 1 summarizes host-balanced retrieval performance with 95% bootstrap confidence intervals using 5,000 resamples. Gram-positive results are shown in Table S2.

**Table 1.** Gram-negative host retrieval performance (host-balanced metrics; 95% bootstrap CIs; best in bold, second-best underlined).

| Model | MRR | Hit@1 | Hit@5 | Hit@10 |
| --- | --- | --- | --- | --- |
| BLASTN | 0.3002 (0.2750–0.3182) | 0.2172 (0.1922–0.2394) | 0.3808 (0.3477–0.3990) | 0.4589 (0.4265–0.4813) |
| VirHostMatcher | <u>0.3207</u> (0.2936–0.3365) | <u>0.2315</u> (0.2027–0.2476) | 0.3902 (0.3599–0.4138) | 0.4929 (0.4652–0.5204) |
| WIsH | 0.2479 (0.2256–0.2627) | 0.1695 (0.1463–0.1858) | 0.3020 (0.2730–0.3212) | 0.4245 (0.3947–0.4474) |
| PHIST | 0.2675 (0.2396–0.2834) | 0.2185 (0.1920–0.2375) | 0.3121 (0.2766–0.3270) | 0.3458 (0.3065–0.3572) |
| Evo2 | 0.3106 (0.2840–0.3232) | 0.1939 (0.1681–0.2095) | <u>0.4368</u> (0.3982–0.4504) | <u>0.5541</u> (0.5248–0.5770) |
| BLASTN+VirHostMatcher+PHIST+Evo2 | **0.3679** (0.3359–0.3785) | **0.2685** (0.2333–0.2804) | **0.4689** (0.4292–0.4847) | **0.5847** (0.5472–0.6025) |

`Evo2` provided the best high-recall retrieval among single methods. On the Gram-negative test cohort, Hit@5 was 0.4368, and Hit@10 was 0.5541 across 308 candidate host species. Its host-balanced MRR was 0.3106 (second best) and within the CI of `VirHostMatcher` (MRR=0.3207), which also achieved the highest species-level Hit@1 (0.2315).

These results suggested a tradeoff at the species level. `Evo2` often ranked the recorded host within the top 10, but was not optimal at placing it first. At higher taxonomic ranks, the task becomes coarser, and genus and family labels capture broader host relatedness. In this case, `Evo2` captured this coarser signal with Hit@1 rising to 0.434 at the genus level and to 0.516 at the family level (Fig 4). It outperformed all single baselines at these ranks. We also report unweighted metrics for all methods in Table S3.

**Fig 4.**
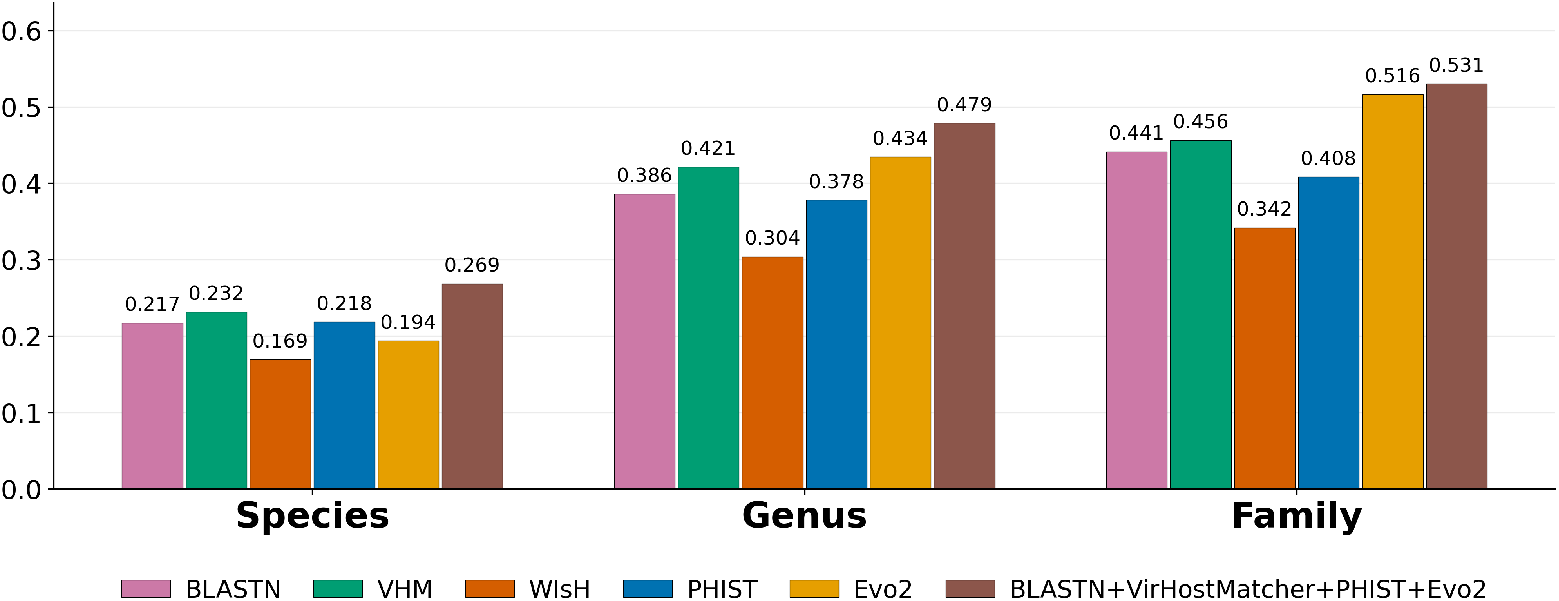
Species, genus, and family top-1 accuracy. Host-balanced Hit@1 is defined as the fraction of phages where the top-ranked host is correct. Values are shown for species, genus, and family on the Gram-negative test cohort. For genus and family, we map each species to its genus or family. The prediction is correct if it matches any recorded host for that phage.

### Unsupervised fusion outperforms single methods

In the previous section, we showed that no single method dominates across the host-balanced MRR and Hit@k. These results suggested that different methods captured complementary signals (e.g., local homology, global composition, or embedding-based similarity). To integrate these signals without training on phage–host labels, we aggregated ranked host lists using reciprocal rank fusion (RRF).

Fig 5 shows the tradeoff between rank quality (MRR) and high-recall retrieval (Hit@10). The results for single methods are shown as stars, and the results for fusions are shown as circles. For each fusion size *n*, we selected one *n*-way fusion using the Gram-positive validation MRR and broke ties using Hit@10. Diamonds mark the selected fusion for each *n*. These are `VirHostMatcher+Evo2` (2-way), `BLASTN+VirHostMatcher+Evo2` (3-way), `BLASTN+VirHostMatcher+PHIST+Evo2` (4-way), and the 5-way fusion.

**Fig 5.**
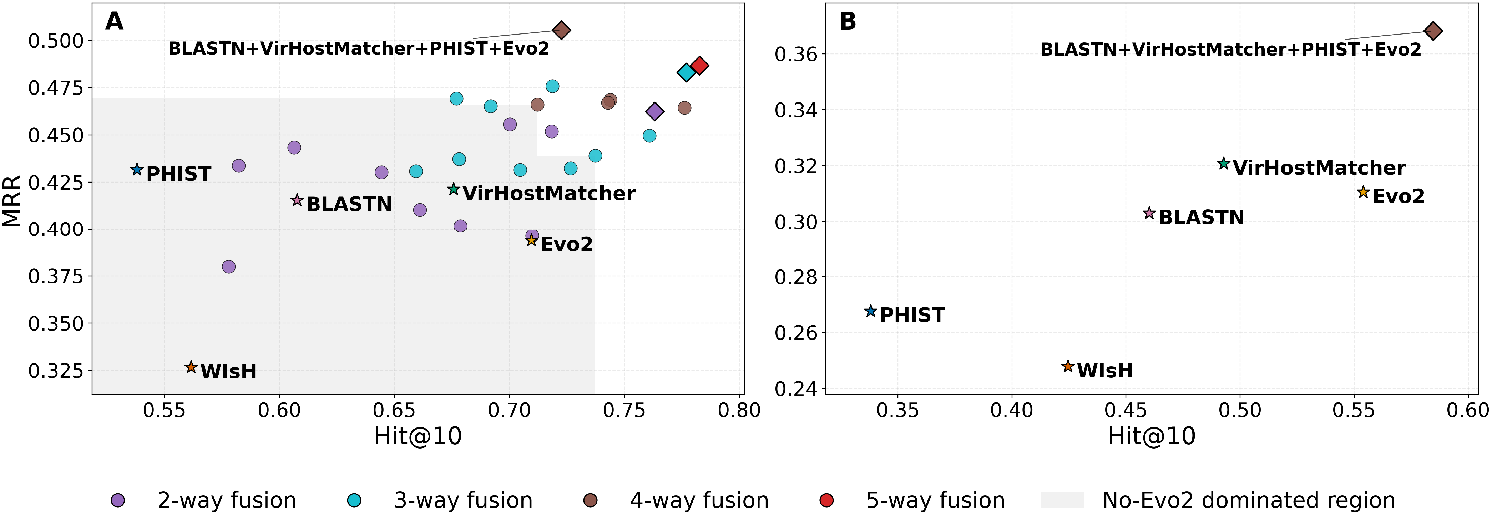
Fusion expands the MRR–Hit@10 frontier. Host-balanced mean reciprocal rank (MRR) is plotted against Hit@10. (A) Gram-positive validation. (B) Gram-negative test. Stars are single methods. Circles are fusions. Circle color shows the number of fused methods. Diamonds mark the validation-selected best *n*-way fusion. The shaded region shows performance without `Evo2`.

On the validation set (Fig 5A; Table S4), moving from the best 3-way to the best 4-way fusion increased MRR by 4.6% (0.483 to 0.505) at a 6.9% cost in Hit@10 (0.777 to 0.723), while the 5-way fusion improved both MRR and Hit@10 by less than 1% beyond the best 3-way. The gray region in Fig 5A highlights the MRR–Hit@10 combinations achievable without `Evo2`. The figure shows that adding `Evo2` expands the attainable MRR–Hit@10 frontier. RRF is label-free and straightforward to apply once the rankings for each of the components are available. Full results for all 1–5-way combinations are in Table S4 (Gram-positive) and Table S5 (Gram-negative).

Based on the Gram-positive validation MRR, we selected the 4-way fusion `BLASTN+VirHostMatcher+PHIST+Evo2` and evaluated this fixed configuration on the Gram-negative test cohort (Fig 5B; Table 1; Table S5). This fusion improved all four species-level metrics, achieving MRR=0.3679, Hit@1=0.2685, Hit@5=0.4689, and Hit@10=0.5847. It also improved genus Hit@1 to 0.479 and family Hit@1 to 0.531 (Fig 4).

### Phage genome length strongly modulates host-prediction performance

We expect genome length to alter which host-range signals were detectable. Short genomes provided less sequence context, whereas longer genomes were expected to better support homology-based signals. We tested these length effects by stratifying the Gram-negative test cohort by genome length (Fig. 6). We used multiple bins in the figure. For a compact summary, we grouped adjacent bins into four length ranges.

**Fig 6.**
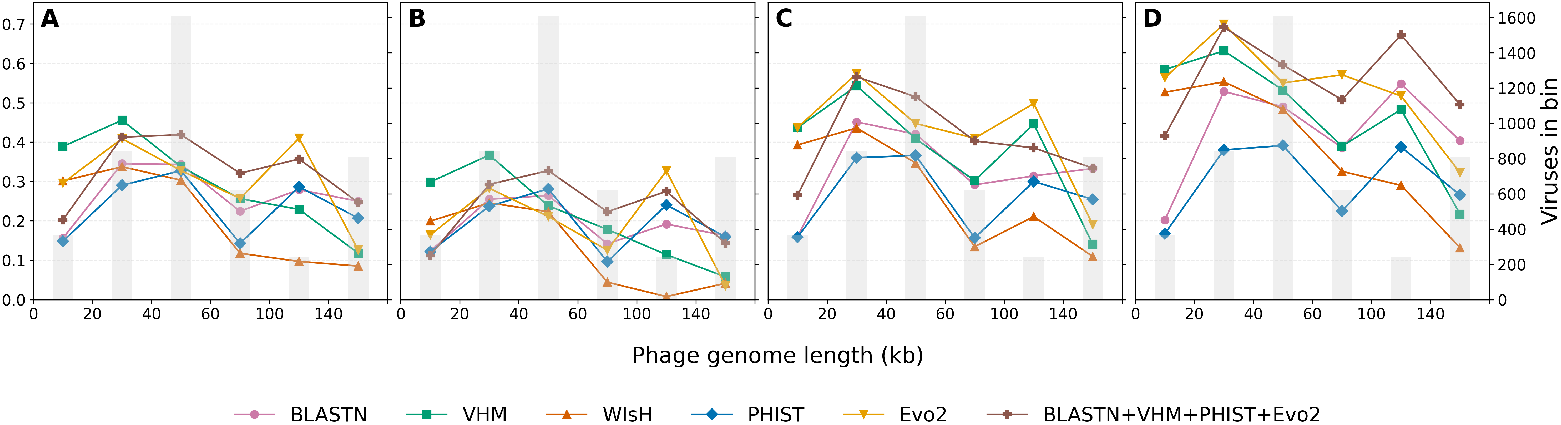
Performance versus phage genome length. Host-balanced MRR and Hit@k on the Gram-negative test cohort, stratified by phage genome length. Lines show single methods and the validation-selected fusion; gray bars indicate the number of phages in each length bin.

For short phage genomes (0–40 kb), `VirHostMatcher` gave the best MRR and Hit@1. In this range, MRR ranged from 0.389 to 0.455, and Hit@1 ranged from 0.299 to 0.367. For intermediate phage genomes (40–100 kb), `Evo2` was consistently the strongest high-recall retriever, achieving the best Hit@5 and Hit@10, even when it did not maximize Hit@1. For 100–140 kb genomes, `Evo2` led MRR, Hit@1, and Hit@5. For very large genomes (*>*140 kb), `BLASTN` dominated all metrics with MRR=0.250 and Hit@10=0.403. `Evo2` dropped to MRR = 0.126. Taken together, these results showed that phage genome size determined prediction strategy.

### Host prediction performance is highly dependent on host clades

Next, we asked whether method performance was consistent across taxa. Host clades varied in the number of phage–host interactions, affecting the sample size of each clade-level estimate. For each cohort, we considered all host taxa present in the dataset from phylum to genus. For each bacterial taxon, we grouped the phages whose recorded host species belonged to that taxon and computed species-level metrics by ranking against the full candidate host database for that cohort. We reported the method(s) with the highest MRR for each taxon. Fig 7 displays the best-performing method across the host taxonomy tree, from phylum to genus, for both cohorts.

**Fig 7.**
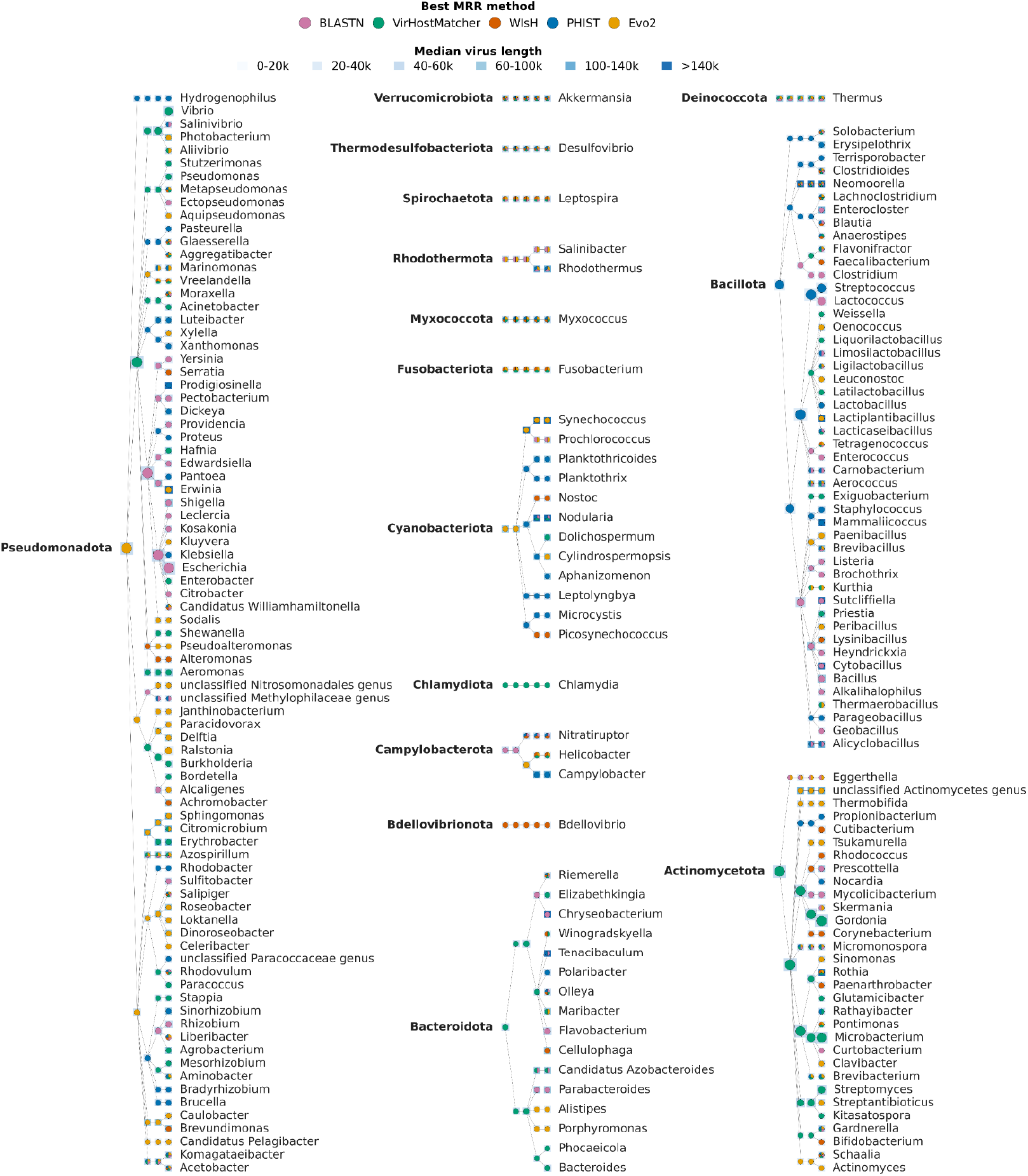
Best-performing method varies across host clades. Host taxonomy trees for the Gram-negative test cohort (left and middle) and Gram-positive validation cohort (right). For each host taxon (phylum to genus), we compute species-level metrics on the associated phages and rank against the full cohort host database. Node color indicates the method with the highest host-balanced mean reciprocal rank (MRR). Ties are shown as pie slices. The halo encodes median phage genome length. Length bins match Fig 6.

In the combined Gram-negative (left/middle) and Gram-positive (right) host taxonomy trees (Fig 7), `Evo2` frequently attained the highest MRR (gold circles) across many major phyla, but different baselines dominated several well-known phyla. This pattern suggested that method performance was strongly clade-specific, and that different lineages were best predicted by different genomic signals. Even within the same phylum, the best-performing method varied across lower taxonomic ranks.

When the analysis was restricted to clades with the longest median phage genomes, `Evo2` was the method that most frequently attained the highest MRR in both cohorts (Table S6). It was also the only method to do so at the order level, for Synechococcales (Cyanobacteriota) and Sphingomonadales (Pseudomonadota), and this advantage extended consistently down to the genus level in both lineages. Across the *Actinomycetes* lineage, `Evo2` led at all taxonomic levels. Furthermore, `Evo2` attained the highest MRR in several genera with median genome lengths exceeding 140 kb, even though `BLASTN` dominated that range across all phages (Fig 6). This divergence indicates that clade-specific genomic signals can override overall length-dependent trends. indicates that clade-specific genomic signals can override overall length-dependent trends.

### Mobile genetic elements modulate the advantage of local homology methods

Our previous results showed that method performance depended on genome length and host clade. We next asked whether mobile genetic elements (MGEs) in the host genome modulated method performance. MGEs are expected to create direct sequence overlap between phages and hosts favoring alignment and exact *k*-mer matching. Insertion sequences (ISs), on the other hand, may increase genomic mosaicism, which may weaken genome-wide compositional signals [9, 22, 37]. We therefore focused on prophages and insertion sequences, both of which are common MGEs involved in horizontal gene transfer [37, 38] We stratified Gram-negative phages by their host MGE nucleotide coverage. Coverage is defined as the fraction of host nucleotides annotated as MGEs. We evaluated host-balanced retrieval metrics within each bin. We also show results for the *validation-selected* fusion (`BLASTN+VirHostMatcher+PHIST+Evo2`) in these stratifications.

#### Integrated prophage coverage predicts when local homology methods gain an advantage

We annotated integrated prophage regions in each bacterial host genome using VIBRANT and computed the fraction of host nucleotides covered by prophages [39]. In Fig 8, we binned each query phage by the prophage coverage of its recorded host (0%, 0–1%, …, ≥4%) and evaluated retrieval against the full Gram-negative candidate host database.

**Fig 8.**
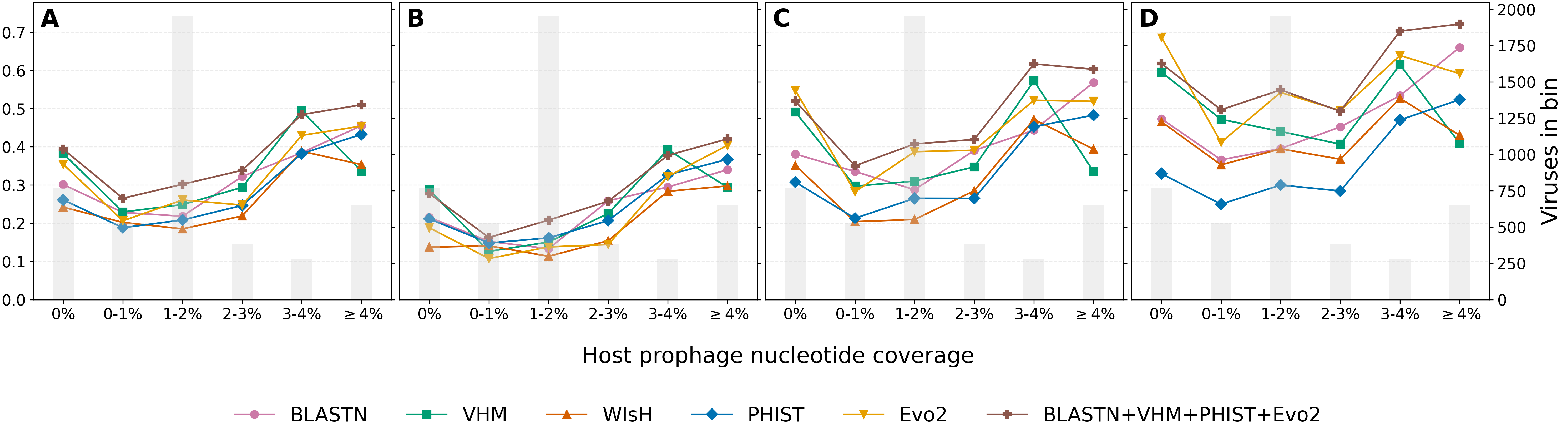
Performance versus host prophage coverage. Panels show host-balanced (A) mean reciprocal rank (MRR), (B) Hit@1, (C) Hit@5, (D) Hit@10 on the Gram-negative test cohort. Phages are binned by integrated prophage nucleotide coverage in the recorded host genome. Prophage regions are annotated with VIBRANT. Gray bars show the number of phages in each bin.

Since some bins contain few phages, differences in method performance across bins need to be interpreted with caution. We observed two patterns. First, higher prophage coverage favored methods based on local matches. These included `BLASTN` and `PHIST`. The fusion curve often matched the best single method. In the ≥4% bin, fusion reached MRR = 0.51 and Hit@10 = 0.72 (Fig 8 A,D), indicating that fusion captured complementary signals and did not collapse to an average performer. Second, at low prophage coverage, `Evo2` remained a strong high-recall retriever. In the 0% bin, its Hit@10 was 0.69. Fusion improved MRR and Hit@1, but its Hit@10 was only 0.62 (Fig 8).

#### Insertion sequence burden induces non-monotonic performance across methods

We next quantified ISs using ISEScan [40]. Hosts were annotated and binned by IS coverage (0–1%, …, ≥3% in Fig 9). ISs are short, often repeat-associated transposable sequences that can increase the abundance of repeated short motifs and introduce shared fragments across distant genomes [38, 40].

**Fig 9.**
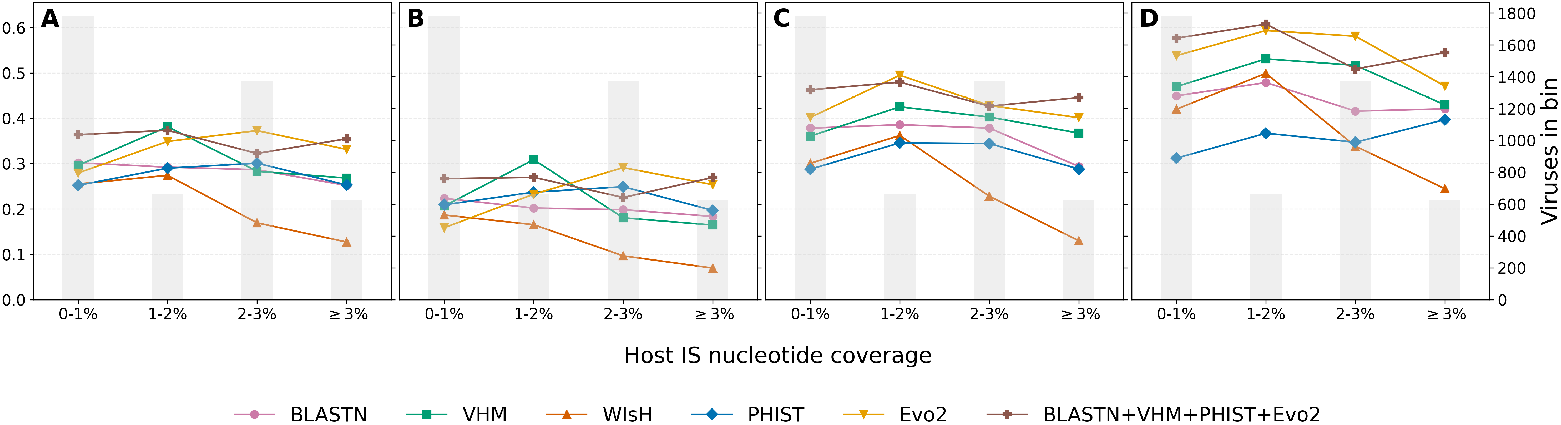
Performance versus host insertion-sequence coverage. Panels show host-balanced (A) mean reciprocal rank (MRR), (B) Hit@1, (C) Hit@5, (D) Hit@10 on the Gram-negative test cohort. Phages are binned by insertion-sequence nucleotide coverage in the recorded host genome. IS elements are annotated with ISEScan. Gray bars show the number of phages in each bin.

Across IS-coverage bins on the Gram-negative cohort, `PHIST` gained recall as IS burden increased, with Hit@10 rising from 0.312 to 0.397, but its MRR and Hit@1 did not improve. `BLASTN` showed similarly limited gains in MRR and Hit@1. This pattern suggested that IS-rich genomes introduce repeated, non-specific matches shared across many hosts rather than stronger host-specific signal. In contrast, `Evo2` remained strong in IS-rich bins. In the 2–3% bin, it achieved the highest MRR and Hit@1. In the ≥3% bin, it achieved the highest Hit@1, and its MRR was close to the best method. This result suggested that the embeddings captured host signal beyond repeated IS fragments (Fig 9). For `Evo2`, Hit@10 ranged from 0.581 to 0.594 in the 1–3% bins and dropped to 0.471 in the ≥3% bin. `WIsH` also degraded sharply in the IS-rich bin. Its MRR was 0.127 at ≥3%.

The validation-selected fusion (`BLASTN+VirHostMatcher+PHIST+Evo2`) was robust across bins and performed well in the most IS-rich bin. In the ≥3% bin, MRR was 0.355 and Hit@10 was 0.545. However, fusion dipped in the 2–3% bin, where `Evo2` alone achieved higher recall. In that bin, Hit@10 was 0.581 for `Evo2` and 0.509 for fusion (Fig 9).

Taken together, these results indicate scenario-dependent method performance on the Gram-negative held-out test cohort. When host genomes were enriched with integrated prophages or ISs, alignment and *k*-mer methods benefited from abundant local-homology evidence, and a fixed fusion could leverage this complementarity to improve overall ranking robustness—especially in the highest-coverage bins. When MGE coverage was low, informative signal was less likely to come from direct matches and more likely to reflect broader genome properties, where `Evo2` remained effective as a top-*k* candidate generator.

## Discussion

Computational phage–host prediction remains a difficult problem in microbiome studies. In this work, we have implemented a new unsupervised host prediction framework for query phages using `Evo2` genomic embeddings. Our approach uses host-balanced metrics: MRR and Hit@k (k=1, 5 and 10) to avoid biases introduced by long tail distributions of hosts in data bases. We benchmarked our method against methods based on alignment (`BLASTN`), oligonucleotide composition (`VHM`, `WIsH`), and shared *k*-mer matches (`PHIST`).

On the Gram-negative test cohort, `Evo2` was the single best method for high-recall retrieval, achieving Hit@5 = 0.4368 and Hit@10 = 0.5541, and was second best for MRR (0.3106). However, it did not maximize species-level Hit@1 (0.1939). At the genus and family levels, `Evo2` outperformed all single baselines, with Hit@1 reaching 0.434 at genus and 0.516 at family. These results suggest that `Evo2` captures information complementary to sequence identity, *k*-mer frequency, and nucleotide composition, and suggest that `Evo2` is most useful as a candidate generator, often placing the recorded host near the top of the ranked list.

We have also proposed a Reciprocal Rank Fusion (RRF) that merges ranked lists without requiring phage–host labels. Our work shows that adding `Evo2` to the baseline tool set improves the performance of any combination and that the selected 4-way fusion (`BLASTN+VirHostMatcher+PHIST+Evo2`), which reached MRR = 0.3679 and Hit@10 = 0.5847, outperforms all other methods at the species level. It also improved genus accuracy to 0.479 and family accuracy 0.531. These results support that prediction of phage-host biology requires other factors not captured by current analysis methods. We explored some of these factors here: phage genome length, clade distribution, prophage and insertion sequence coverage.

We showed that phage genome length strongly modulates the performance of the methods. Prior reports on phage–host prediction show that alignment-free composition/Markov signals degrade on shorter viral contigs, while homology and shared-*k*-mer signals strengthen as longer sequence becomes available [13–15, 23]. These studies, however, varied contig length down to 1–10 kb, whereas our stratification used complete-genome lengths with coarser bins. We find that for short phage genomes (0–40 kb), `VirHostMatcher` led MRR and Hit@1, while `Evo2` placed third in MRR and Hit@1 but led Hit@5. For intermediate lengths (40–100 kb), fusion led MRR and Hit@1, while `Evo2` achieved the best Hit@5 and Hit@10. In the 100–140 kb range, `Evo2` led MRR, Hit@1, and Hit@5. For the longest genomes (*>*140 kb), `BLASTN` led MRR, `PHIST` led Hit@1, and fusion led Hit@10. Even in this longest-genome bin, overall performance was lower than in mid-length bins (Fig 6). This pattern was consistent with long genomes aligning well and suggested that whole-genome pooling could dilute signal in this regime. Across length bins, `Evo2` was most consistently strong in recall-oriented retrieval, finishing first in Hit@5 in four of six bins, but it rarely maximized Hit@1 outside the 100–140 kb range. Our results on length dependence argue, therefore, for hybrid pipelines rather than a single default method.

Performance also varied across host clades, even within the same phylum. Because related bacteria often inhabit similar environments, these clade-level differences may partly reflect environment-dependent variation in the genomic signals available for prediction [13, 23]. In *Pseudomonadota*, `BLASTN` often led in *Enterobacterales* genera such as *Escherichia* and *Citrobacter*, which are clinically relevant and extensively sampled in Virus-Host DB [41]. `VirHostMatcher` often led for genera such as *Pseudomonas* and *Acinetobacter*. These bacteria are commonly encountered in soil, water, and hospital-associated environments, suggesting a strong role for genome-wide compositional similarity in these clades [42, 43]. Agriculturally relevant hosts showed mixed performance across tools. For example, `Evo2` led for *Erwinia* (a well-known plant-associated lineage), while other plant-associated genera such as *Agrobacterium* and *Xanthomonas* favored `VirHostMatcher` and `PHIST`, respectively [44–46]. Within *Bacillota* (Gram-positive), `PHIST` led for several clinically important genera, including *Staphylococcus* and *Streptococcus*. These clade-level switches show that host context should guide tool choice and fusion. Fig 7 provides a practical view for clade-aware prioritization that different clades are better captured by different kinds of host-related information. In some clades local homology is most informative, whereas in others compositional or embedding-based similarity is more informative. These signals may reflect phage–host co-evolution. When coarse priors are available, they can guide which methods to run first. In regions of the host taxonomy where no single method consistently dominates across neighboring taxa, fusion is especially beneficial as it combines complementary signals.

Host mobile genetic elements helped explain variation in method performance. High prophage coverage (≥4%) was associated with stronger performance of alignment and *k*-mer exact-match methods, consistent with local sequence overlap. At very low prophage coverage (0%), `Evo2` retained the highest Hit@5 and Hit@10, `VirHostMatcher` led Hit@1, while fusion achieved the highest MRR. One possible explanation for why the recorded host may be ranked lower on the fused list when prophage coverage is low is that direct-match methods tend to disagree more than in high-coverage hosts at the top ranks, and RRF favors hosts that are placed near the top by multiple methods. Insertion sequence coverage affected prediction methods differently from prophage coverage. Whereas prophage regions provided shared sequence that favored alignment and *k*-mer methods, IS elements increase repetitive content throughout the host genome, which may homogenize oligonucleotide profiles across unrelated hosts and reduce their discriminative power (`VirHostMatcher` and `WIsH`). `Evo2` embeddings were more robust to IS burden, achieving the highest MRR at 2–3% IS coverage. These observations suggest that the type and abundance of mobile genetic elements in candidate hosts is an important consideration when selecting or combining prediction methods.

Direct-evidence signals remain important, but they are often sparse. We evaluated CRISPR spacer matching with SpacePHARER, but it identified matches for only 14.6% of phages. SpacePHARER also did not produce scores for all phage–host pairs in our benchmark. For this reason, we excluded it from the main comparisons. In practice, sparse evidence could still rerank a shortlist produced by methods that score all candidates.

We additionally benchmarked three machine learning models on the Gram-negative test cohort: `DeepHost` [26], `CL4PHI` [29], and `VirHostMatcher-Net` [25]. `DeepHost` is a CNN classifier trained on known phage–host interactions. Its species model covers 40 of our 308 test hosts and its genus model covers 189. The species model reached a species Hit@1 of 0.093 and the genus model reached genus Hit@1 of 0.197. `CL4PHI` embeds phages and hosts into a shared space via contrastive learning and is capable of scoring arbitrary host genomes. Evaluated against all 308 hosts, it reached a species Hit@1 of 0.129. `VirHostMatcher-Net` combines CRISPR spacer matches, alignment, and oligonucleotide composition within a Markov random field over 62,493 prokaryotic reference genomes. It covered 260 of our 308 hosts and reached a species Hit@1 of 0.154. All three models fell below the four baselines and `Evo2`, largely because their training host sets covered only a fraction of the 308 test species. The true host is unknown at prediction time, so it is difficult to assess whether a given query falls within a model’s training domain. Unsupervised methods, by contrast, produce dense rankings over all candidate hosts without relying on interaction labels.

Our evaluation uses a closed-world retrieval setup. Each query phage ranks hosts within a fixed candidate database built from complete genome assemblies of Virus-Host DB host species. If the true host species or the relevant strain is missing, no method can return a correct label. Because we collapsed strain labels to species, strain-level specificity is also hidden. Expanding the candidate host database will enable evaluation in more realistic open-world settings. A further limitation is that cosine similarity and RRF produce ranking scores rather than the calibrated probabilities. The pipeline therefore always returns a ranked host list, even for out-of-scope queries such as viruses with eukaryotic hosts. Developing confidence measures that support abstention is an important direction for practical deployment.

As shown by the performance reduction in IS-rich hosts, equal-weight fusion can inherit biases from component methods when individual signals degrade unevenly. Fusion strategies that adapt to host genomic context could mitigate this effect. Embedding extraction remains computationally intensive. On four NVIDIA A5000 GPUs, embedding a bacterial host genome took ∼25 minutes and a phage genome ∼2 minutes.

Overall, frozen `Evo2` embeddings provide a broad and useful host-range signal. Combined with simple rank fusion, they support robust host shortlisting and ranking across diverse host taxa. Our stratified analyses offer practical guidance for building hybrid pipelines in new datasets.

## Methods

### Virus-Host DB cohorts and preprocessing

We used the *Virus-Host Database (Virus-Host DB)*, a curated repository that links viruses and their hosts using NCBI taxonomy identifiers (TaxIDs) [34]. Virus-Host DB integrates virus–host information from RefSeq, GenBank, UniProt, and ViralZone, supplemented by manual literature curation [47–50]. We used the October 2024 release of Virus-Host DB (release219), which contained 36,470 virus–host pairs with both prokaryotic and eukaryotic hosts.

For this study, we formed two bacterial cohorts. The Gram-negative cohort contained 4,600 phage–host interactions. It contained 4,490 unique phages and 308 host species. Among these phages, 4,477 were DNA viruses and 13 were RNA viruses. The Gram-positive cohort contained 3,538 interactions. It contained 3,514 unique phages and 209 host species. All phages in the Gram-positive cohort were DNA viruses. We assigned Gram status using phylum-level heuristics. We treated the Gram-positive cohort as the validation set for embedding and fusion choices and reserved the Gram-negative cohort as a held-out test set for all final reported performance.

These cohorts were filtered subsets of Virus-Host DB (see criteria below), not the full set of Gram-labeled entries. Virus-Host DB reports hosts at multiple taxonomic ranks. We retained interactions whose host was annotated at the species or strain level, but we collapsed all host labels to species TaxIDs in NCBI taxonomy for evaluation. We further required that each candidate bacterial host species had an available *complete* genome assembly from NCBI, and we used one representative assembly per host species to define the candidate host database. We retrieved phage genome sequences from Virus-Host DB directly.

### Embeddings

`Evo2` is pretrained on 9.3 trillion DNA base pairs spanning all domains of life. It employs a StripedHyena 2 multi-hybrid architecture with model scales from 1B to 40B parameters and a context window of up to one million tokens. Without task-specific fine-tuning, `Evo2` embeddings support downstream biological tasks. Recent work uses `Evo2` for genome modeling and design [32], including generative design of viable bacteriophage genomes in *E. coli* [33]. The `Evo2` model computes a stack of blocks, each producing a hidden-state representation (embedding) of the input sequence. Prior work in `Evo2` shows that different blocks encode different levels and types of information, and downstream tasks can prefer intermediate blocks [32]. In this work, we kept `Evo2-7B` frozen, represented each host and phage genome using pooled hidden states from a selected intermediate block, and performed unsupervised host retrieval by similarity.

#### Whole-genome embedding extraction

To obtain a fixed-dimensional embedding for a full genome *g*, we tiled the sequence with fixed-length windows, embedded each window, and pooled token embeddings across the genome. We used 8,192-nt windows and 25% overlap (stride 6,144 nt). Although `Evo2` supports longer contexts, we chose this window length to match `Evo2`’s base pretraining context length and to keep whole-genome embedding computationally feasible on our hardware [32]. The 25% overlap supplied boundary context and reduced windowing artifacts. Because `Evo2` is decoder-only (left-to-right), token embeddings depended only on upstream context. Thus, for tokens in the overlap, the earlier window provided more left context than the later window. To avoid double-counting, we pooled only the non-overlapping suffix (6,144 nt) from each window after the first. We then collected retained token embeddings from all windows and computed a single mean over all tokens to obtain a 4,096-dimensional vector.

#### Embedding-space normalization

Before cosine retrieval, we normalized embeddings by (i) per-dimension *z*-scoring and then (ii) *ℓ*_2_ normalization. The *z*-scoring statistics (*µ, σ*) were estimated from a reference set ℛ_*c*_. We evaluated three unsupervised choices for ℛ_*c*_: (i) hosts only, ℛ_*c*_ = ℋ_*c*_; (ii) hosts plus the external phage bank, ℛ_*c*_ = ℋ_*c*_ ∪ ℬ; and (iii) external phage bank only, ℛ_*c*_ = ℬ. We denote the candidate host genome database by ℋ_*c*_ for each cohort *c*, and ℬ is defined below. We selected the reference-set choice on the Gram-positive cohort and then fixed it for all downstream analyses.

#### External phage bank ℬ

The phage bank ℬ was constructed from Virus-Host DB by removing (i) all query phages used in the Gram-positive and Gram-negative cohorts and (ii) any phage whose species-level host label appeared in either candidate host database, ensuring that ℬ was disjoint from both evaluation phages and candidate host taxa. The resulting bank contained |ℬ| = 1,500 phages (750 infecting Gram-positive hosts and 750 infecting Gram-negative hosts). Host annotations were used only to construct a disjoint reference set, and the normalization used the embeddings only.

Formally, let *e*(*g*) ∈ ℝ^4096^ denote the `Evo2` genome embedding of a genome *g* under the fixed extraction procedure, and let *e*(*g*)_*j*_ denote its *j*-th coordinate. We computed per-dimension empirical mean and standard deviation over the reference set:

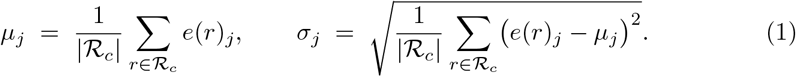

For any host or phage embedding *x* = *e*(*g*), we applied a reference-set-based *z*-scoring followed by *ℓ*_2_ normalization:

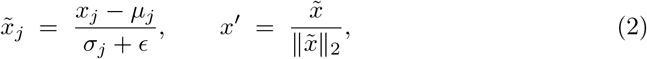

where a small constant *ϵ* is added for numerical stability. When ℛ_*c*_ included ℋ_*c*_, (*µ, σ*) were computed per cohort; when ℛ_*c*_ = ℬ, the same (*µ, σ*) were shared across cohorts.

#### Cosine retrieval

Given a query phage *v* and candidate host *h*, we computed cosine similarity *S*(*v, h*) = *v*′^⊤^*h*′ and ranked candidate hosts for each phage by decreasing *S*(*v, h*).

#### Block choices

We treated the `Evo2` block used for *e*(*g*) and the normalization reference set as validation choices selected on the Gram-positive cohort and then fixed for all downstream analyses.

### Retrieval and Rank

For each cohort *c*, we denote the query phage genomes by 𝒱_*c*_. Each *v* ∈ 𝒱_*c*_ is a *query phage* whose host is retrieved by ranking all candidate host *species* genomes *h* ∈ ℋ_*c*_. Virus-Host DB records host TaxIDs per phage *v* at the strain/species level. We first map any strain-level TaxID to its parent species and define the species-level recorded host set 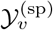. We report host-balanced metrics at multiple taxonomic ranks *r* ∈ {sp, genus, family}. Let *π*_*r*_(·) map a *species* TaxID to its rank-*r* TaxID (with *π*_sp_ the identity), and define the deduplicated true label set at rank *r*:

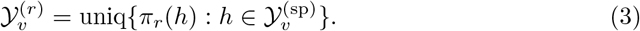

A prediction is counted as correct at rank *r* if the top-ranked *species* host maps to any element of 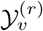 Accordingly, we define the best (lowest) retrieved rank among all candidate hosts whose mapped label matches:

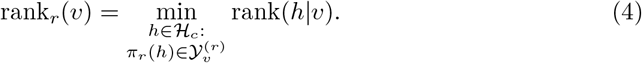

Because host frequencies in Virus-Host DB are strongly long-tailed (Fig 2), we assess retrieval using host-balanced Mean Reciprocal Rank (MRR) and Hit@k. For each evaluation rank *r*, we define the label frequency over *phages*:

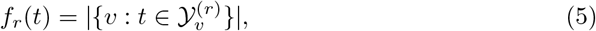

and introduce a weighting term that inversely scales by label frequency:

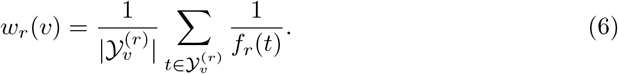

Then the host-balanced MRR at rank *r* is:

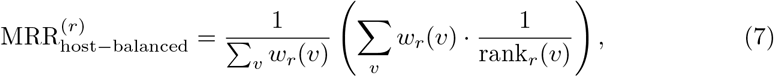

and the host-balanced Hit@k is:

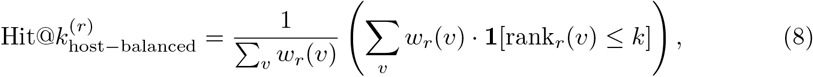

where **1**[·] is the indicator function.

The MRR reflects the average inverse rank of the true label, emphasizing how close to the top a correct label appears. Hit@k measures the proportion of queries for which at least one correct label appears among the top-*k* ranked predictions. Host-balanced metrics prevent common hosts (e.g., *E. coli*) from dominating evaluation and instead weight each host label equally at the chosen rank *r*.

### Baseline phage–host prediction models

To benchmark our approach, we compared against four widely used unsupervised phage–host prediction tools. In all cases, each method was given the same set of viral genomes and candidate host genomes. For a labeled phage, each method produced a similarity score over candidate hosts. For tools that could return sparse outputs (e.g., PHIST and BLASTN), we filled missing pairs with zero similarity. If a query had zero similarity to all candidates, we treated that method as providing no discriminative ranking (all hosts tied).

#### BLASTN

`BLASTN` provides an alignment-based baseline that detects direct nucleotide similarity between each viral–bacterial genome pair. For each phage–host pair, we consider the best alignment between the two genomes and use its bitscore as the similarity score. Hosts with strong (higher bitscore) alignment to a phage are ranked as more likely hosts [12]. We ran `blastn` from the BLAST+ package (v2.17.0) with `-task blastn` against a host-genome database built with `makeblastdb -dbtype nucl`, using default parameters.

#### VirHostMatcher (VHM)

`VirHostMatcher` is an alignment-free method that compares the overall oligonucleotide composition of viral and host genomes. It summarizes each genome by its k-mer frequency profile and uses the 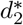 dissimilarity metric to quantify profile dissimilarity. Hosts whose k-mer composition is more similar to that of a phage (lower 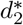 dissimilarity) are ranked as more likely hosts [13]. We used the default `d2*` settings provided in the VirHostMatcher (v1.0.0) implementation and ranked hosts by increasing 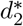.

#### WIsH

`WIsH` trains a separate Markov model of nucleotide composition for each candidate host genome. It learns the characteristic sequence patterns of each host and evaluates each phage by how well the host-specific model fits the phage genome. A likelihood score per phage–host pair is produced. Hosts whose model assigns higher likelihood to a viral genome are ranked higher as candidate hosts [14]. We ran WIsH (v1.1) with default parameters, training host models with `WIsH -c build` and scoring phages with `WIsH -c predict`.

#### PHIST

`PHIST` takes a *k*-mer-based approach that counts shared exact *k*-mers between phage and host genomes. Phages that share more *k*-mers with a host genome are more likely to infect that host. Hosts sharing more *k*-mers with a phage receive higher ranks [15]. We ran PHIST (v1.2.1) with default parameters (including *k* = 25).

### Fusion of host rankings

The unsupervised predictors captured complementary host-range signals. To integrate them without training on phage–host labels, we used reciprocal rank fusion (RRF). For each phage *v*, each method *m* produced a ranking over the same candidate host set, which yielded ranks rank_*m*_(*h*|*v*) for host *h*. RRF aggregated these ranked lists by assigning each candidate host a fused score

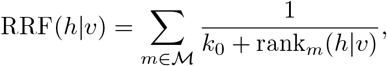

where ℳ is a subset of methods being fused and *k*_0_ controls how quickly contributions decay with rank. We used *k*_0_ = 60 for all methods following standard RRF practice and then re-ranked hosts by descending RRF(*h*|*v*). The fused output is a single ranked list of candidate hosts per phage.

### Host mobile genetic element annotation

We annotated integrated prophage regions in each bacterial host genome using VIBRANT and computed the fraction of host nucleotides covered by prophages [39]. We quantified insertion sequences (ISs) using ISEScan and computed IS nucleotide coverage per host genome [40]. For stratified analyses, each query phage was assigned the prophage and IS coverage of its recorded host species, and performance was evaluated within coverage bins while ranking against the full candidate host database.

## Data Availability Statement

Code for data processing, Evo2 embedding extraction, and evaluation is available at https://github.com/Arsuaga-Vazquez-Lab/Evo2-Host-Prediction.git; phage genomes and virus–host associations were downloaded from Virus-Host DB (https://www.genome.jp/virushostdb/), and candidate bacterial host genomes were downloaded from NCBI using the NCBI Datasets command-line tools (https://github.com/ncbi/datasets).

## Acknowledgments

JA thanks Prof. A. Bhatt for receiving him in her laboratory during JA’s sabbatical.

## Supporting information

**S1 Fig. Whole-genome Evo2 embedding**. Each genome is tiled into 8,192-nt windows with 25% overlap (stride 6,144 nt). Each window is embedded with `Evo2-7B` at a chosen block; to avoid double-counting overlap tokens, only the non-overlapping suffix is retained after the first window. Token embeddings are then averaged across all retained positions to form a single 4,096-dimensional genome vector.

**S2 Fig. Reference-set normalization workflow**. (A) Construct an external reference set (phage bank). It is disjoint from evaluation phages and host taxa. (B) Estimate per-dimension mean and standard deviation on the reference set. Apply *z*-score normalization and *ℓ*_2_ normalization. (C) Rank candidate hosts for each phage by cosine similarity in the normalized embedding space. (D) Normalization improves host-balanced mean reciprocal rank (MRR) and Hit@k relative to raw embeddings.

**S1 Table. Reference-set normalization ablation for Evo2 retrieval**. Host-balanced MRR and Hit@k for Evo2 embedding-based host retrieval under four normalization choices: no reference-set normalization, hosts only, hosts plus the external phage bank, and external phage bank only. For normalized conditions, embeddings are z-scored per dimension using statistics estimated on the reference set and then *ℓ*_2_-normalized prior to cosine-similarity ranking.

**S2 Table. Gram-positive host retrieval benchmark**. Gram-positive cohort results reported as host-balanced MRR and Hit@k with 95% bootstrap confidence intervals (5,000 resamples over viruses), evaluated by ranking each query phage against the full Gram-positive candidate host database.

**S3 Table. Standard retrieval metrics**. Unweighted (“standard”) retrieval metrics at species level (MRR and Hit@k) and Hit@1 at genus and family ranks. Main-text results use host-balanced metrics to correct for long-tailed host frequencies.

**S4 Table. Fusion ablation on the Gram-positive cohort**. Host-balanced MRR and Hit@k for all 1–5-way reciprocal rank fusion (RRF) combinations over the five component methods (BLASTN, VirHostMatcher, PHIST, WIsH, and Evo2). Checkmarks indicate which methods are included; underline and bold highlight within-*n* and overall best configurations, respectively.

**S5 Table. Fusion ablation on the Gram-negative cohort**. Host-balanced MRR and Hit@k for all 1–5-way reciprocal rank fusion (RRF) combinations over the five component methods (BLASTN, VirHostMatcher, PHIST, WIsH, and Evo2). Checkmarks indicate which methods are included; underline and bold highlight within-*n* and overall best configurations, respectively.

**S6 Table. Long-genome clade winners by method**. Counts of long-genome host clades for which each method attains the highest host-balanced MRR; ties are counted for all tied methods and results are stratified by cohort and taxonomic level.

## Notes

### Competing Interest Statement

The authors have declared no competing interest.

